# The Hidden Prior: Variance Constraints Under Data Augmentation

**DOI:** 10.64898/2026.08.31.748288

**Authors:** E. G. Cooch, D. I. MacKenzie, J. A. Royle

## Abstract

Data augmentation is now a standard device across capture–recapture and occupancy analysis: adding a fixed number M of all-zero encounter histories replaces a model of unknown dimension with one of fixed dimension. Although M is often treated as a computational tuning choice, it also specifies a finite superpopulation and hence a binomial support constraint on the number of undetected individuals. In a Bayesian implementation that constraint appears as an induced prior; in a likelihood implementation it is the same finite-support assumption reached by another route. That the Bernoulli specification for the inclusion indicators induces a binomial prior on abundance is established (Schofield & Barker 2014); our concern is what that choice costs in estimated uncertainty. We develop the argument using a simple closed-population abundance estimation problem. We show that augmented occupancy and Huggins conditional-likelihood analyses give numerically identical point estimates of 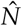 once M is sufficiently large. Their uncertainty estimates, however, need not agree. We distinguish two sources of discrepancy. First, when M is small relative to the number of undetected individuals, the finite binomial ceiling truncates the likelihood or posterior and suppresses uncertainty. Second, once that ceiling no longer binds, Taylor-series (Delta-method) approximations still understate variance, because the quantity of interest is a strongly non-linear function of the estimated parameters and local linearization does not reproduce its curvature. Gauss–Hermite quadrature on the unconstrained logit scale recovers much of the shortfall and approaches the MCMC posterior benchmark, though a small residual remains that does not close as M grows, reflecting the distinction between asymptotic likelihood theory and finite-sample Bayesian inference. Neither mechanism is peculiar to abundance estimation: the first follows from the augmented representation itself, the second from any derived quantity that is a non-linear function of estimated parameters. We close with framework-specific guidance for choosing M.

---

Data augmentation through the addition of a known number of latent all-zero encounter histories is now commonly used in mark–recapture analysis, to account for those individuals that were present in the sampled population, but never caught. Since its introduction as a general method for multinomial models with an unknown index (Royle, Dorazio & Link 2007), the technique has been used in a variety of capture–recapture analyses (Royle *et al*. 2009; Gardner *et al*. 2010; Royle & Dorazio 2012), and in the hierarchical formulations that are increasingly used in quantitative wildlife monitoring (Royle, Converse & Link 2012; Kéry & Schaub 2012).

Across these diverse applications, data augmentation is primarily valued for its computational elegance: it replaces an unknown-dimensional problem with a fixed-dimensional one, allowing for inference in frameworks where marginalizing out *N* is analytically intractable (Royle, Dorazio & Link 2007; Royle *et al*. 2009; Gardner *et al*. 2010; Royle & Dorazio 2012). However, this ubiquity comes with a persistent, under-examined technical challenge: in every one of these formulations – from simple closed-population models to more complex open-population and spatial capture-recapture (SCR) models the choice of the augmentation size *M* is not merely a tuning parameter for Markov chain Monte Carlo (MCMC) efficiency (Royle, Converse & Link 2012; Kéry & Schaub 2012). As we will demonstrate, it is the specification of an implicit prior, and one that carries systematic consequences for uncertainty quantification that persist even as sampling frameworks grow in complexity (Link 2013).

For most practitioners, data augmentation is first encountered – and is still most often applied – within a Bayesian framework. For example, consider the estimation of animal abundance, *N*, using encounter data from a closed-population mark–recapture study. In a standard Bayesian framework, treating population size as an unknown parameter creates an immediate technical problem: the dimension of the model changes with *N*. If there are *N* individuals in the population, then there are *N* encounter histories, *N* individual latent states, *N* individual random effects if they are included, and so on. If *N* changes during the MCMC, the number of things being sampled changes with it.

In principle, this can be handled using Reversible-Jump MCMC (Green 1995), but that approach is mathematically and computationally burdensome. Data augmentation avoids this problem by replacing an unknown-dimensional problem with a fixed-dimensional one, by augmenting the observed encounter histories with a fixed (known) number *M* of all-zero histories, giving a fixed superpopulation size *N*_max_ = *D* + *M*, where *D* denotes the number of distinct individuals encountered at least once.^1^ Then, for each of the *N*_max_ potential individuals, we introduce a latent indicator variable, say *z*_*i*_, where *z*_*i*_ = 1 if individual *i* is a member of the population, and *z*_*i*_ = 0 otherwise. The actual population size is then a simple derived quantity,

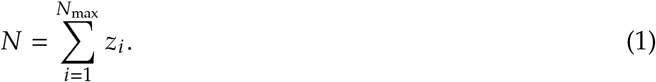

Thus, the MCMC always samples the same number of latent variables, regardless of the realized value of *N*. The abundance problem has been converted into an ‘occupancy-style’ problem: among the fixed set of possible individuals, which ones are actually present in the population? This re-expression is especially useful in Bayesian hierarchical models. Once the individual membership indicators *z*_*i*_ have been introduced, additional structure can be added in a relatively natural way. Individual heterogeneity in encounter probability can be modeled using individual random effects, temporal variation can be handled through occasion-level random effects, and spatial capture-recapture models can include latent activity centers for the augmented individuals. Conceptually, the model simply says: first decide whether a potential individual exists, and conditional on its existence, model its encounter process. Individuals with *z*_*i*_ = 0 are structural zeros; individuals with *z*_*i*_ = 1 are members of the population that may or may not have been detected.

Within this Bayesian paradigm, *M* is generally introduced as a *computational* quantity. Each augmented individual enters the model as a latent binary variable that the sampler must update, so computational cost scales linearly with *M*. The practitioner is accordingly advised to balance two competing pressures: *M* must be large enough to comfortably exceed the (unknown) number of undetected individuals (*f*_0_), but not so large as to make an already expensive sampler intolerably slow. In this view, choosing *M* is a matter of ‘tuning’ – a trade-off between run time and the size of the latent state space.

Our goal in this short paper is to demonstrate that this computational trade-off, while seemingly innocuous, potentially obscures an important statistical consideration. The choice of a specific, finite *M* is not only a decision about ‘computational run time’; it is the specification of an implicit prior. By fixing *M* we are in fact imposing a finite upper bound on the number of possible unobserved individuals, and in the augmented representation the number of occupied all-zero histories is then treated as Binomial with mean *Mψ*_*c*_, where *ψ*_*c*_ is the conditional probability that an undetected individual is in fact present. This Binomial assumption is the ‘hidden’ prior of our title. (In the standard Bayesian construction the same structure is explicit: assigning a uniform prior to *ψ* with *z*_*i*_ *∼* Bernoulli(*ψ*) induces a discrete uniform prior on *N* over {0, 1, … , *N*_max_}.)

The Binomial structure follows from the particular prior placed on the inclusion indicators, not from data augmentation as such. Schofield & Barker (2014) showed that the Bernoulli specification *z*_*i*_*∼*Bernoulli (*ψ*) induces exactly this Binomial prior on *N*, and gave alternative specifications for *z* under which *N* is Poisson (up to truncation at *M*), removing the Binomial constraint while retaining the fixed-dimension representation. Because the Bernoulli form is the one used in nearly all applied work, it is the case we treat throughout. Our contribution is not the observation that the induced prior is Binomial, which is established, but the demonstration that this choice has systematic and quantifiable consequences for the estimated variance of 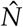.

Although the ‘data augmentation’ device is most closely associated with Bayesian inference, its underlying mechanics are entirely independent of whether you are a Bayesian or a frequentist, and can just as readily be applied within a maximum-likelihood framework. In fact, it might be argued that using ‘data augmentation’ potentially provides substantial practical advantages in a frequentist framework: it leverages widely available, highly optimized occupancy software such as the R packages **unmarked** (Fiske & Chandler 2011), **RPresence** (MacKenzie & Hines 2023), and the **RMark** (Laake 2013) interface to **MARK** (White & Burnham 1999). This makes it quite straightforward to incorporate complex spatial or individual covariates, and makes the full suite of likelihood-based tools – such as AIC-based model selection, multi-model inference, and model averaging (Burnham & Anderson 2002) – directly accessible without the need to formulate bespoke likelihood functions for an unknown *N*. Bayesian and likelihood-based analyses differ in how inference proceeds, but the augmented data structure – and the logical equivalence it encodes – is common to both. We illustrate this using a familiar example concerning estimating animal abundance, and demonstrate that equivalence not merely conceptually but mechanically.

## 1 Equivalence of Point Estimates Between Methods

Consider a closed-population mark–recapture study where live encounter data are collected across *K* sampling occasions. For each individual, we record a binary encounter history: 1 if detected on that occasion, 0 otherwise. A zero entry unambiguously indicates a false negative – an animal present in the population but not detected on that sampling occasion.

For example, the following encounter history data were generated by numerical simulation of sampling from a closed population, where *N* = 135, *p* = 0.5, where *p* is the probability of encounter, across *K* = 3 sampling occasions (left-hand side; Table 1). Since *p* < 1, there is a non-zero probability that an animal is missed on a particular sampling occasion. Estimation of abundance from such data involves summing the number *known* alive in the population over the sampling period (*D*; individuals seen at least once – this represents the minimum population size) with the number *estimated* to be alive, but not observed (*f*_0_). For the simulated encounter data for example, the number of individuals encountered at least once is *D* = 116. This is an observed count, and not a statistical estimate. The statistical challenge in estimating population size is in fact estimating the number of individuals alive and in the population that were not encountered, 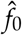.

**Table 1:**
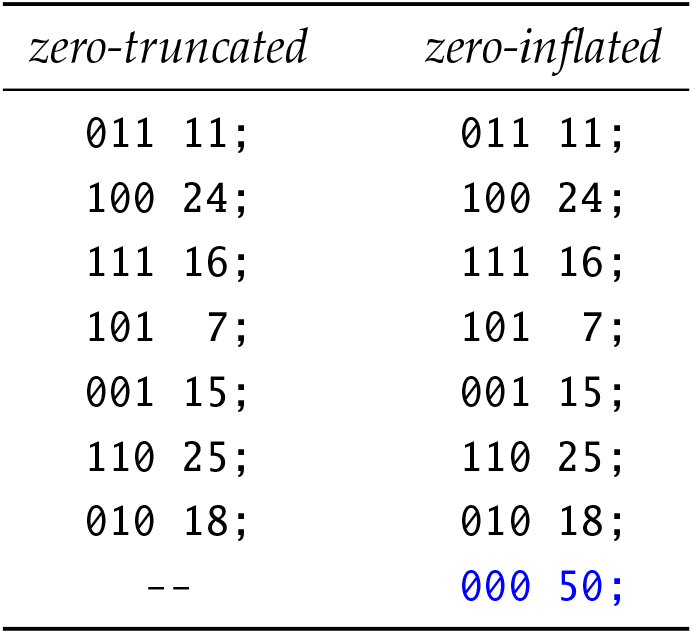
Comparison of zero-truncated encounter data (left-hand column) with version of the data that has been zero-inflated (‘augmented’) by the addition of M = 50 histories consisting of all zeros, ‘000’ (right-hand column). zero-truncated zero-inflated.

There are at least 3 general approaches to estimating 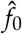. The first is the traditional approach, which maximizes a multinomial likelihood based on the frequencies of the observed encounter histories. This likelihood underlies the various classic closed-population abundance estimators (Otis *et al*. 1978). In this approach, *f*_0_ is a parameter in the data likelihood, and abundance is estimated as a derived parameter by summing 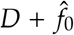. Another approach conditions on the animals that are actually observed (i.e., the analysis is restricted to the *D* individuals encountered at least once; Huggins 1989, 1991). Because *f*_0_ is unobserved, it drops out of the likelihood equations, and the model estimates only the initial capture *p* and recapture *c* probabilities. Using these estimates, the probability that any single individual in the population is caught at least once during the entire study (typically denoted as *p*^*\**^) is derived, and abundance 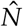 is estimated as the ratio 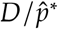. The number of unobserved individuals *f*_0_ is derived simply as 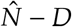. Both the unconditional (Otis *et al*. 1978) and conditional (Huggins 1989, 1991) approaches use ‘typical’ encounter data, where only individuals encountered at least once are included.

A third, alternative approach, as noted earlier, is to (in effect) treat the problem of abundance estimation as a ‘patch occupancy’ problem (MacKenzie *et al*. 2002, 2006). This approach relies on data augmentation (Royle, Dorazio & Link 2007), where the original encounter data are ‘augmented’ by adding a fixed number of ‘missing individuals’, such that the maximum size of the population is known (Table 1), and the problem is recast as a patch occupancy problem. The observed encounter history data (Table 1, left) contains only individuals detected at least once – a zero-truncated sample. The underlying binomial encounter process generates both detections and non-detections, but only the former appear in the field dataset. This truncation is a consequence of the sampling process, not an artifact of data collection.

We are going to manually *augment* the data set, by adding a known number *M* (where *M* > *f*_0_) of ‘000’ histories to the file. Augmenting the data set is the opposite of truncating it. And, by adding ‘000’ histories to the encounter file, we create a ‘*zero-inflated*’ set of histories. For example, in the right-most column of Table 1, we have augmented the observed encounter histories by adding *M* = 50 histories consisting of all zeros, ‘000’ (the right-hand column of the table). For the augmented data, *N*_max_ = *D* + *M* = 116 + 50 = 166.

The key conceptual link between DA for capture-recapture and site occupancy models involves regarding that each individual in the closed population as (in effect) a ‘patch’. An animal that is not in the population is thus an ‘individual patch’ that is simply unoccupied. Consider again the zero-inflated histories in Table 1. We have *M* = 50 ‘patches’ in the encounter file which may be ‘unoccupied’ (i.e., have a ‘000’ history). The question we consider is – what proportion of them are, in fact, really occupied? In other words, if we consider these ‘patches’ as individuals, our question then becomes ‘how many of the individuals with ‘000’ encounter histories really *occur* in the population?’. How many are likely to represent individuals that are alive in the population? In order to answer this question, we need an estimate of the *proportion* of these ‘undetected individuals’ that are occupied.

It might seem reasonable to assume that we could estimate this number simply as 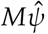, using the estimate 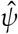 from a standard single-season occupancy analysis applied to our encounter data. However, we need to be a bit careful here. In our present situation, we have a fixed, *finite* number of patches (i.e., the total number of patches is the finite sum *D* + *M*). Unlike most occupancy studies, we are not dealing with a random sample from a large, potentially infinite population. In that case, the occupancy parameter *ψ* is the *probability* that a randomly selected site in that infinite population is occupied. What we need here, then, is an estimate of the *proportion* of the finite number of *M* individuals which are ‘occupied’. This is strictly equivalent to estimating *f*_0_ in the classic approach to abundance estimation from zero-truncated data.

To determine the proportion of the *M* patches with ‘000’ encounter histories that are likely to be occupied, we need an estimate of the *conditional* probability that a given patch is in fact occupied, given that it was not detected as such. Following MacKenzie *et al*. (2002, 2006), this conditional inclusion probability, *ψ*_*c*_, is given by Bayes’ theorem as

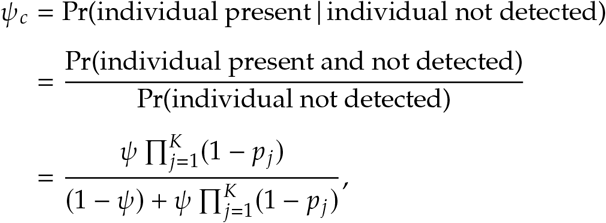

which, if the encounter probability *p* is constant over time (which we assumed in our simulation of the encounter histories), can be simplified to

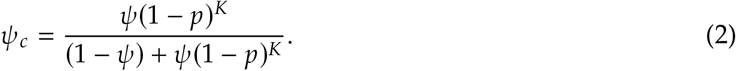

Using this conditional probability, *ψ*_*c*_, we can estimate the proportion of the *M* individuals which are ‘occupied’ simply as the product 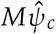. Thus, our derived estimate of abundance would be

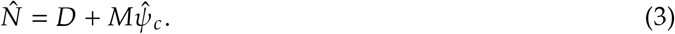

Operationally, then:

1. Take the standard zero-truncated encounter history file and augment it with a known number *M* of ‘000’ histories.
2. Analyze the augmented data set as a single-season occupancy model, yielding 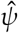 and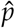.
3. Derive 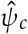 from eqn. 2, the probability that an augmented individual is present given that it was never detected.
4. Estimate abundance as 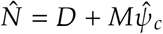 (eqn. 3).

The key practical question is: how many all-zero histories should be added to the encounter file (i.e., what value to use for *M*)? One must add at least as many all-zero histories as there are true, undetected individuals alive in the population—that is, *M* ≥ *f*_0_. If *f*_0_ is unknown, the practitioner must either estimate it from preliminary data or choose *M* conservatively to ensure this condition is met. In the simulated example, the true abundance is *N* = 135, so *f*_0_ = (*N*− *D*) = (135 −116) = 19, and therefore *M* = 50 satisfies the requirement.

### 1.1 Numerical demonstration

We demonstrate this equivalence numerically. For the simulated data with *D* = 116 observed individuals, we augmented by adding *M* = 50 all-zero histories and fit a single-season occupancy model ({*ψ*· , *p* (.)}) to recover parameter estimates. Using program **MARK**, the Huggins conditional likelihood approach on the original zero-truncated data yielded 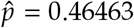 and 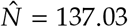 (SE = 6.858).

Applying the augmented-data occupancy approach: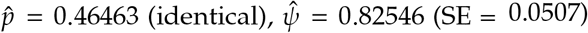, and thus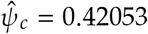. From eqn. 3, 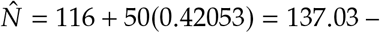 numerically identical to the Huggins estimate. The point estimate of *N* is invariant to *M* for all *M* > *f*_0_ = 19; we verified this by repeating the analysis with *M* = 100, 150, 200 and obtained the same estimate in each case. However, as we demonstrate next, the choice of *M* is consequential for uncertainty quantification.

## 2 The variance trap: binomial ceilings and mathematical loss

Standard occupancy or capture–recapture software generates an asymptotic variance estimate for 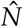. It is helpful to briefly review the derivation of the variance estimator. Consider the joint log-likelihood of the total population size (*N*) and the detection probability (*p*). The likelihood is modeled as a binomial process where *N* individuals are subject to *K* trials. Treating *N* as continuous, the asymptotic variance is found by calculating the Fisher information matrix from the expected values of the negative second derivatives of the log-likelihood.

Inverting the matrix yields the top-left element for:

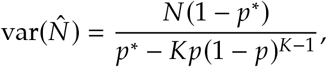

where *p*^*\**^ is the probability of detecting an ‘individual’ at least once after *K* survey occasions (for constant *p, p*^*\**^ = 1 − (1 − *p*)^*K*^).

The critical feature of this estimator is what it assumes about *N*. By treating *N* as a continuous, unbounded quantity, there is no hard upper limit on how many individuals might have gone undetected. Conversely, under data augmentation, an upper-bound *M* is imposed and, if this is not sufficiently large, it will lead to truncation of the likelihood (or posterior distribution). This can lead to bias in the statement of uncertainty.

In the data augmentation framework, the population size *N* is conceptually replaced by the expected number of occupied patches in the augmented sample. Because *s* is the total augmented superpopulation (*s* = *D* +*M*) and *ψ* is the unconditional inclusion probability, the expected population size is *E* [*N*] = *sψ*. By direct analogy to Darroch’s (1958) classical closed-population variance formula, and substituting the augmented-data parameters (*sψ, p*^*\**^) in place of the classical-model parameters, the corresponding finite-sample variance estimator is (assuming constant detection probability and no unsampled patches):

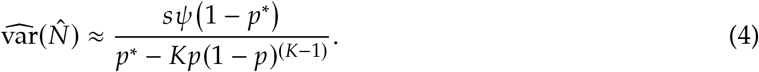

This heuristic connection rests on recognizing that the augmented-data occupancy model and classical closed-population capture–recapture share the same underlying binomial sampling structure; the variance formulae are thus parallel under the parameter substitution *N* → *sψ*.

Because 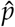 (and thus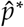) is invariant to *M*, and 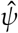 is a simple function of *M*, we expect that the variance from this estimator should be invariant to *M*.

This is exactly what is observed. For example, given 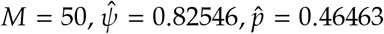 (such that *p*^*\**^ = 0.84655), *s* = (116 + 50) = 166, with *K* = 3 sampling ‘surveys’, then

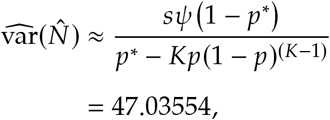

which gives 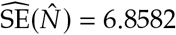. This value (6.8582) is what is reported by **MARK** for model *M*_0_,^2^ using the Huggins closed-model likelihood.

### 2.1 Derivation of the variance from first principles

It is instructive to consider the derivation of the variance estimator from a different perspective. Let *z*_*i*_ be the binary occupancy state for site *i*. If we observed the true state for every site (i.e., if *p* = 1) then the number of occupied sites *x* is the quantity

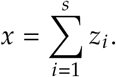

However, in practice, *p* < 1, and thus the occupancy state is not known for some sites. In that case, the estimator of *x* is

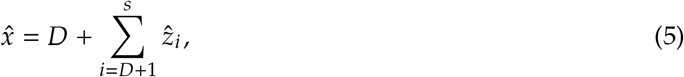

where *s* is the number of sampling units, and *D* is the number of units (patches, individuals) in sample *s* where the species (or, in the present context, the individual) is detected. For convenience, we’ll let *n* = (*s* − *D*).

Because the expected value of the second term in eqn. 5 is *nψ*_*c*_, we can re-write eqn. 5 as

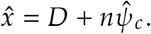

In other words, for the situation where all possible sites in the finite sample are visited, *x* is estimated as the number of occupied sites in which the species was detected plus the estimated occupancy status of each site where no detections occurred. Note that this is equivalent to eqn. 3, where *x* ≡ *N* and *n* ≡ *M*.

Thus, we could re-write eqn. 3 as

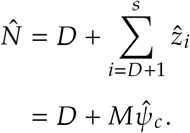

From eqn. 3, 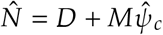. Because *D* is an observed count, and not an estimated parameter, then 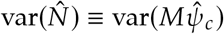.

Using conditional expectations, we derive an estimator for var 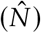from first principles (recalling that *n* ≡ *M*) as

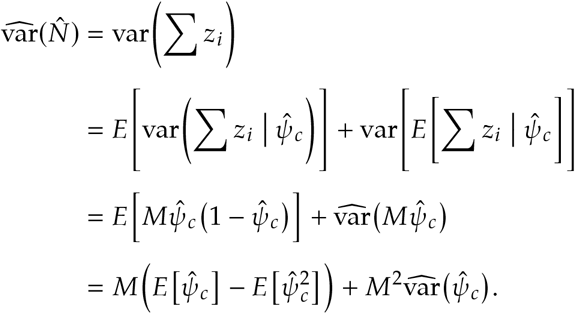

Recalling that

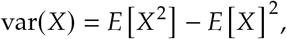

such that

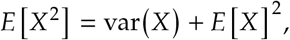

then continuing from above

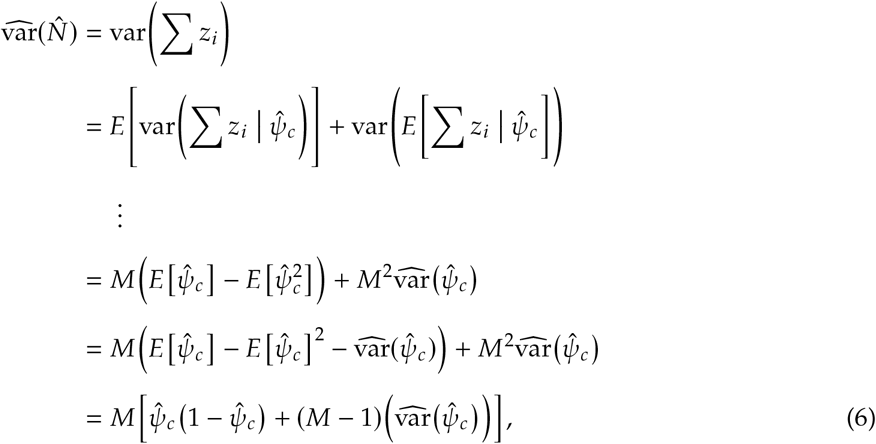

where 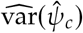 is approximated by application of the Delta method to eqn. 2. For our present example, using the *M* = 50 augmented data set, 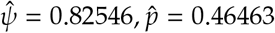 (such that *p*^*\**^ = 0.84655), *s* = 116 50 = 166, and *K* = 3 sampling occasions (surveys). From eqn. 2, 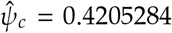, with 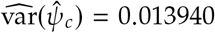 (Addendum 1).

Then from eqn. 6,

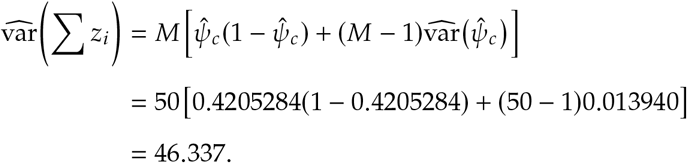

Thus, the estimated SE is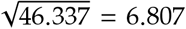, which is close to – but not exactly the same as – the reported value from the Huggins abundance estimation using program **MARK** (6.858), or calculated using eqn. 4. Why the difference?

### 2.2 Reconciling the two estimators: the binomial ceiling

The two variance estimators are not unrelated; in fact, they differ by a single, interpretable term. We can reconcile the Darroch asymptotic MLE estimator with our conditional derivation from first principles as follows. We first expand our derived estimator eqn. 6:

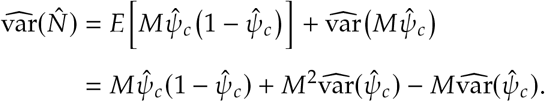

By dropping the term –*M*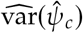 , we effectively remove the upper bound imposed by a finite *M*. This reduced expression amounts to assuming an infinite augmented population (*M*→∞), shifting the estimator’s premise from a bounded Binomial system to an unbounded one. Doing so recovers the asymptotic variance. This modified formulation, which we will refer to here as eqn. 7, converges structurally to the standard Darroch estimator introduced earlier:

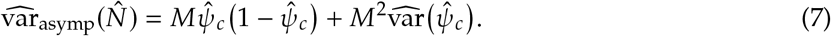

If we use this expression, substituting in estimates derived earlier, we get

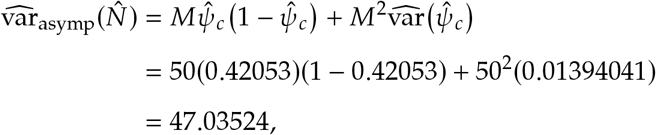

which is nearly identical to the variance estimate that comes out of **MARK**, or from eqn. 4, above.

Now, compare this back to our strict derivation from first principles in eqn. 6, which we expand here as eqn. 8:

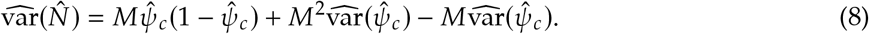

Written this way, we see that the first two terms on the RHS of eqn. 8 are identical to eqn. 7, the asymptotic formulation. The only difference is that our conditional derivation eqn. 8 requires the subtraction of *M* 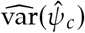. This term results in the estimated variance using eqn. 8 being smaller than the asymptotic estimate from eqn. 7 (and by extension, the variance estimate from **MARK**). Because the difference between the two estimators is exactly *M*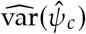, then as *M* ↑ , the magnitude of this product drops (as *M* ,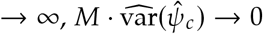). This is exactly what you see if you increase *M* – the difference between the strictly derived variance from eqn. 8 and the asymptotic value reported by **MARK** or eqn. 7 gets smaller and smaller (for the sample data, for *M* ≥ 300, the difference is almost zero).

So, which of the estimators for 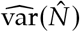 (i.e., with or without the finite population correction term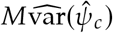) is ‘correct’? The formula for the variance based on conditional expectations eqn. 8 intrinsically depends on the mechanics of data augmentation. By choosing a specific, finite value for *M*, we are explicitly imposing a finite upper bound on the number of possible unobserved individuals. In the augmented representation, the number of occupied all-zero histories is treated as Binomial with mean *Mψ*_*c*_. This Binomial assumption is key because it places an upper limit on the potential population size, which artificially constrains the variance.

In contrast, standard closed-population models (and by extension, the asymptotic estimator) do not assume a hard upper bound; they effectively treat the underlying population size as an unbounded random variable. When a small, finite *M* is used, truncation of the likelihood occurs and will systematically underestimate the variance. However, as the number of trials (*M*) increases, the likelihood becomes sufficiently close to zero that the effect of this truncation is negligible.

This convergence explains exactly what is observed as the number of augmented histories is increased: the artificial ceiling is lifted. Therefore, the asymptotic estimator behaves as if no upper limit is imposed, avoiding the artificial variance constraint of a bounded sample.

Thus, data augmentation forces an explicit Binomial assumption that can systematically underestimate your variance if *M* is too small. Jaber *et al*. (2023) make the same point in a spatial-capture setting, framing the augmented size as the binomial trial count on which estimate accuracy depends. This is the hidden prior in its most concrete form: the choice of *M* is the choice of a ceiling, and the ceiling is paid for in variance.

#### 2.2.1 What the correction term is – and what it is not

The −*M* 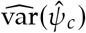 term warrants clarification. Athough it diminishes as *M*→ ∞ and resembles a finite-population correction, it is technically a second-moment correction arising from *estimation* of *ψ*_*c*_ rather than a sampling correction in the classical sense. The within-sample binomial variance component 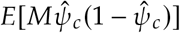 and this estimation-uncertainty term are distinct effects: the former drives the binomial ceiling when *M* < *f*_0_, whereas the latter is a comparatively small correction that vanishes as *M* grows.

A second distinction matters for what follows. The upper-limit just described is a *finite-M* effect: it has the largest influence when *M* is small relative to *f*_0_ – most notably if *M* < *f*_0_. However, even with large *M*, we will show that estimators based on the Delta method will invariably be biased low, because the conditional inclusion probability *ψ*_*c*_ has a heavy right tail that a local, polynomial approximation cannot see. This is not strictly a Binomial-ceiling effect, but rather a property of the approximation.

## 3 The true variance by MCMC

Although the preceding sections demonstrated the mechanics of data augmentation using a maximum-likelihood framework, the technique is most often associated with Bayesian inference. We compare our frequentist results directly against a Bayesian implementation to see how the theoretical variance derivations perform empirically – and to establish an empirical benchmark for the “true” finite-sample variance against which our analytical approximations can be judged.

It is straightforward to fit the same underlying abundance model *M*_0_ in a Bayesian MCMC framework. Following Kéry & Schaub (2012), we implemented the Bayesian model using **rjags** (v. 4.7, linked to JAGS 4.3.2) in **R** (Version 4.6.1; **R** Core Team 2026), executing the MCMC sampler across three parallel chains to ensure convergence. We retained a standard uniform prior on the encounter probability (*p ∼*Uniform (0, 1)) and a uniform prior on the inclusion probability (*ψ ∼*Uniform (0, 1)), and defined the latent inclusion state (*z*_*i*_) for all potential individuals up to *N*_max_ = *D* +*M*. For each augmentation scenario (i.e., different values for *M*), the model was initialized with 1,000 adaptation iterations followed by 2,000 burn-in updates. Posterior distributions were generated from a total of 10,000 iterations per chain. Mean parameter values were calculated over 100 runs of the samplers, at each value of *M*.

A word on the prior for *N* is warranted here, because the choice interacts directly with the quantity we are trying to measure. The flat prior on *ψ* that underlies standard data augmentation is equivalent to a discrete uniform prior on *N* over { 0, 1, … , *N*_max_} , and as *M* grows this approximates a constant prior on the positive integers. Link (2013) showed that the constant prior is not the innocuous default it appears to be: it can yield an improper posterior, and even where the posterior is proper it produces inferences that depend on *M* in ways that are hard to defend. Link recommended the scale prior [*N* ] *∝*1 / *N* instead, which is approximated in practice by replacing *ψ ∼*Uniform (0, 1) with *ψ ∼*Beta (0.001, 1). We therefore ran every augmentation scenario under both specifications. In the present *M*_0_ example – 116 detections, constant *p*, and no individual heterogeneity – the posterior is proper and well behaved under either specification, and the two gave practically indistinguishable results. We accordingly report uniform-prior results below, which are simpler to interpret: the finite augmented ceiling *N*_max_ = *D* + *M* constrains the posterior variance predictably when *M* is small, with the constraint increasingly lifted as *M* increases beyond *f*_0_. Link’s cautions apply with considerably more force in the sparse and heterogeneous settings that motivated them, and nothing here should be read as evidence against them.

Table 2 compares the empirical posterior variance from the MCMC chains against the analytical estimators. For the moment we consider only the first-order column (Delta_1_), derived using the first-order approximation to 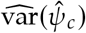 in eqn. 6; the remaining columns are discussed in Section 4. Each row in the table represents the mean of 100 independent MCMC runs: the point estimates 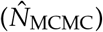 are posterior means, and the variance estimates are the mean of the posterior variances computed over those 100 replications. As *M* increases, point estimates for 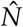 converge, confirming the logical equivalence of the two frameworks (Section 1.1). However, for safely large values of *M* (*M* ≥ 45), the MCMC variance is systematically higher than the analytical variance.

**Table 2:** Comparison of mean empirical MCMC variance (calculated over 100 replications) against standalone maximum-likelihood analytical approximations for 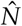 across different augmentation sizes (M). True f_0_ = 19. MCMC point estimates 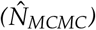 are posterior means; variance estimates are the mean of the posterior variances from each run. The analytical methods are deterministic variance estimators derived from asymptotic likelihood theory: ‘Delta_1_’ represents the standard first-order Taylor series approximation; ‘Delta_2_’ applies a second-order correction to account for the local curvature of the likelihood surface; and ‘GH_Logit_’ performs numerical quadrature on the unconstrained logit space, allowing the symmetric nodes to naturally capture the heavy right-hand tail of the unobserved individuals distribution when back-transformed.

| $M$ | $\hat{N}_{\text{MLE}}$ | $\hat{N}_{\text{MCMC}}$ | $\widehat{\text{var}}_{\text{MCMC}}$ | Delta<sub>1</sub> | Delta<sub>2</sub> | GH<sub>Logit</sub> |
| --- | --- | --- | --- | --- | --- | --- |
| 15.00 | 130.99 | 128.60 | 4.98 | 0.05 | 0.05 | 50.52 |
| 30.00 | 137.03 | 136.14 | 28.58 | 45.67 | 47.29 | 38.77 |
| 45.00 | 137.03 | 137.84 | 48.16 | 46.24 | 47.23 | 42.06 |
| 60.00 | 137.03 | 137.91 | 50.14 | 46.48 | 47.24 | 44.47 |
| 75.00 | 137.03 | 137.90 | 50.61 | 46.61 | 47.26 | 45.71 |
| 90.00 | 137.03 | 137.91 | 50.52 | 46.69 | 47.29 | 46.43 |
| 105.00 | 137.03 | 137.91 | 50.55 | 46.75 | 47.30 | 46.89 |
| 120.00 | 137.03 | 137.92 | 50.52 | 46.79 | 47.32 | 47.19 |
| 135.00 | 137.03 | 137.88 | 50.52 | 46.82 | 47.33 | 47.41 |
| 150.00 | 137.03 | 137.94 | 50.64 | 46.84 | 47.35 | 47.58 |
| 200.00 | 137.03 | 137.91 | 50.49 | 46.90 | 47.37 | 47.91 |
| 250.00 | 137.03 | 137.87 | 50.34 | 46.92 | 47.39 | 48.08 |
| 300.00 | 137.03 | 137.90 | 50.70 | 46.94 | 47.40 | 48.18 |

This discrepancy highlights a limitation in the first-order Delta method on which the analytical variance estimate is based: because the transformation to derive *ψ*_*c*_ places the detection probability *p* in the denominator, the resulting posterior for *N* is asymmetrical with a heavy right-hand tail. The MCMC sampler – a global integration tool – fully captures the mass of this heavy tail, whereas the first-order Delta method is a local, linear approximation that effectively truncates it, leading to an underestimate of the true uncertainty. Fig. 1 illustrates this effect empirically, visualizing how the posterior distribution transitions from a hard-truncated shape at low *M* to a heavy-tailed, right-skewed state as *M* provides sufficient “room” for the estimate to converge.

**Figure 1:**
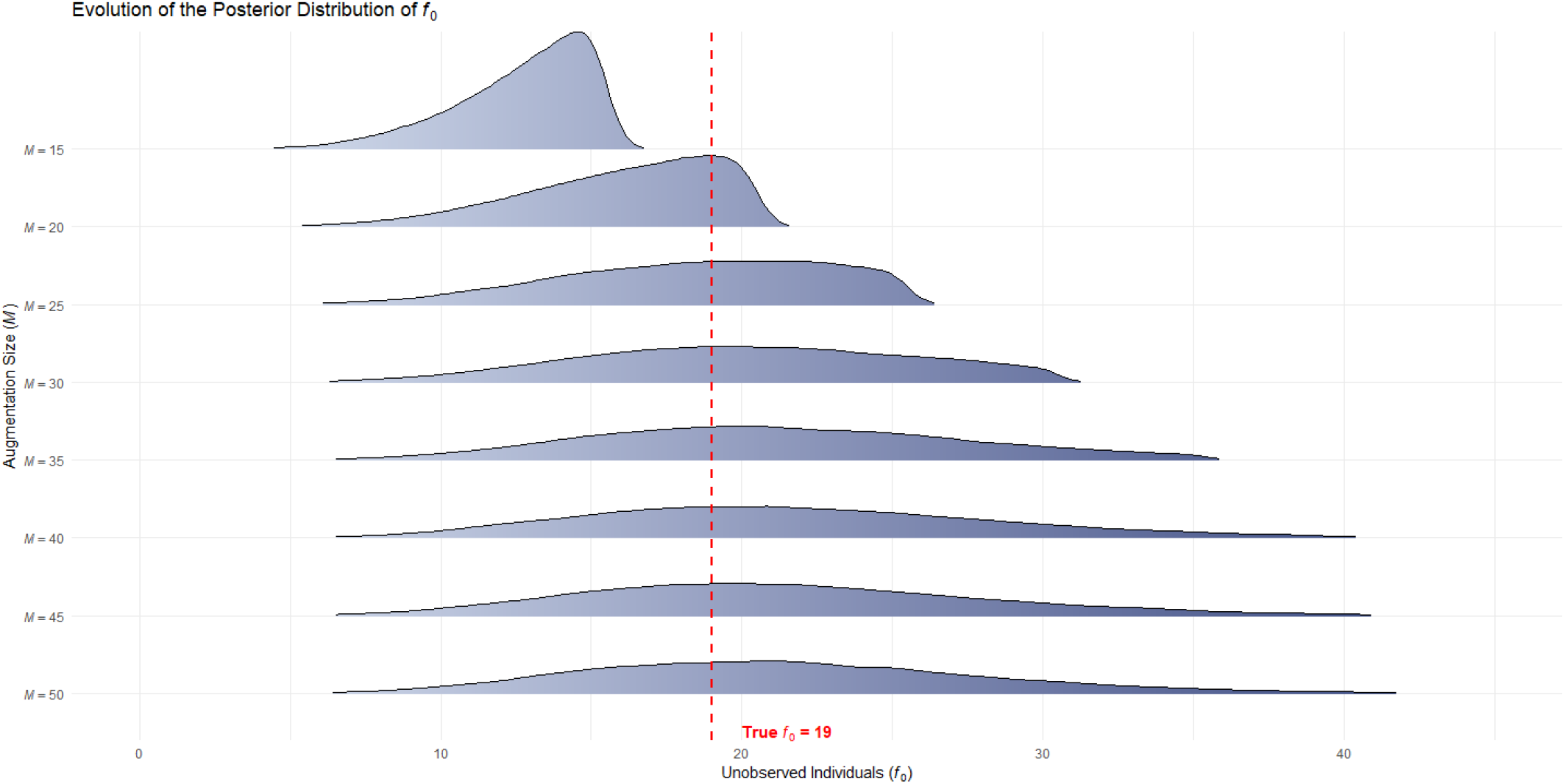
Evolution of the posterior distribution of f_0_. Note how the distribution transitions from a hard-truncated shape at low M to a heavy-tailed, right-skewed state as M provides sufficient “room” for the estimate to converge.

## 4 Improving the estimate of var(*ψ*_*c*_)

If we wanted to stay within a frequentist ML-based framework, there are a variety of options we might consider to improve our estimate of 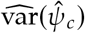. First, we might consider simply increasing the order of the Taylor series approximation (Addendum 2). By accounting for the curvature of the non-linear transformation via the Hessian matrix, the analytical estimate physically expands to capture the right-skewed mass of the distribution (Fig. 1).

However, a Taylor expansion – of any finite order – cannot account for the full volume of a distribution that is strongly asymmetrical. Because the conditional probability formula for *ψ*_*c*_ places the detection probability *p* in the denominator, the true likelihood surface for the number of unobserved individuals (*f*_0_) is asymmetrical, with a strong right skew (Fig. 1). Because the Taylor expansion assumes symmetric (Gaussian) local behavior, it cannot “see” the full volume of this heavy tail. Consequently, it artificially clips the distribution, leading to a systematic underestimation of the true finite-sample variance. Once the augmentation size *M* exceeds the true number of unencountered individuals (*f*_0_), the point estimates are fixed in place. Consequently, the local curvature stabilizes, and Delta estimators will hit an artificial variance ceiling.

Another more flexible approach is based on Gauss-Hermite quadrature (GH; Abramowitz & Stegun 1964). GH performs a *global* integration. As such, if we match the exact distributional assumptions of the asymptotic likelihood theory underlying the data, we can use GH to evaluate the exact transformation function over the true probability mass, so that the GH variance estimate scales naturally with *M*. We matched those assumptions by mapping the standard normal nodes into the logit parameter space (Addendum 3).

Comparisons of 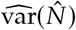 obtained under each treatment of 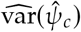 (first- and second-order Delta hods, and GH integration) are presented in Table 2.

First, consider the situation where *M* < *f*_0_ (i.e., *M* = 15) in Table 2. In this situation, the augmented ceiling *N*_max_ = *D* +*M* = 116 +15 = 131 lies *below* the true population size (135). As a consequence, the conditional inclusion probability *ψ*_*c*_ is forced toward 1.0, and the MLE itself is visibly constrained against the boundary (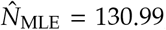, with *ψ*_*c*_≈ 1). This boundary constraint invalidates standard asymptotic maximum-likelihood theory: when the parameter estimate is pinned at the boundary of the parameter space, the usual regularity conditions for asymptotic normality no longer hold (Self & Liang 1987), and the curvature of the likelihood surface at the boundary becomes essentially flat.

Consequently, every variance estimator fails at this design point (*M* = 15), though not in the same way. The asymptotic normal approximation to the likelihood surface is misspecified: the logit-scale covariance matrix describes a distribution whose mass spills well beyond the hard ceiling (Fig. 2). When the logit-scale numerical quadrature (GH_Logit_) integrates across those nodes, it produces an inflated variance (50.52) that is an artifact of integrating over a misspecified approximation rather than a meaningful uncertainty estimate. The Delta estimators collapse to near zero in the same row because they are anchored at an MLE that is pinned against the boundary, where local curvature is essentially flat and standard asymptotic results fail. Both failures – the inflated quadrature estimate and the collapsed Delta estimates – are direct symptoms of the boundary constraint, not of the methods themselves.

**Figure 2:**
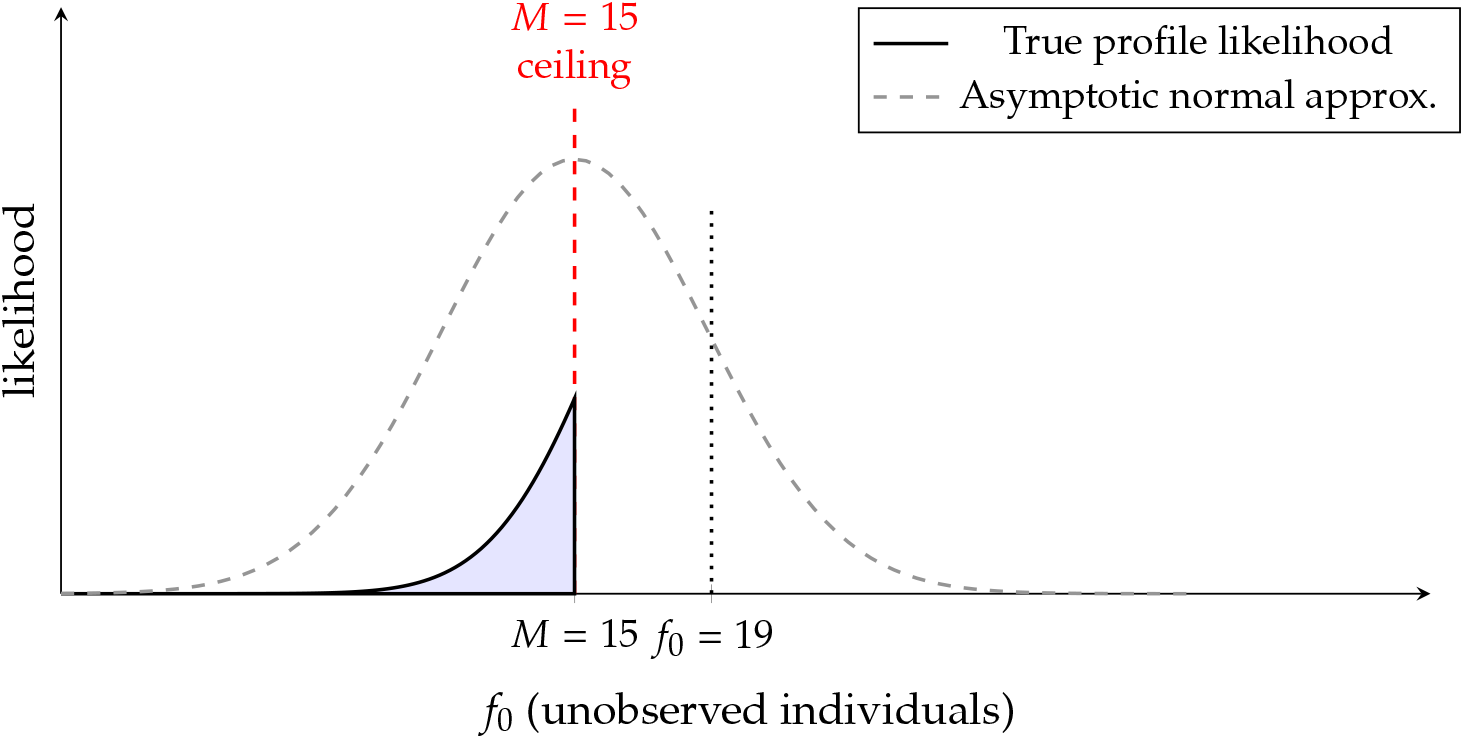
Schematic of likelihood misspecification at the binomial ceiling. The true profile likelihood for f_0_ is maximized at the true value (f_0_ = 19) but is hard-truncated at the augmentation ceiling M = 15, so the estimate is pinned against the boundary and the surface is still rising where it is cut off. The asymptotic normal approximation anchored at that pinned estimate places substantial mass beyond the ceiling, on values of f_0_ the augmented model cannot represent.

These failures are symptoms of *M* < *f*_0_ and are increasingly mitigated as *M* increases beyond *f*_0_. For *M* ≥ 45 (i.e., once *M* is safely larger than *f*_0_), the empirical MCMC variance recovers from the boundary constraint, and the MLE is no longer constrained by the ceiling. At this point, the first- and second-order Delta methods hit their local mathematical ceiling: from *M* = 45 onward both fall short of the empirical MCMC variance, and that shortfall does not close as *M* grows. In marked contrast, the logit-scale numerical quadrature (GH_Logit_) smoothly and monotonically captures the expanding, right-skewed volume of the distribution as *M* grows.

However, it is worth noting that the GH_Logit_ variance remains below the empirical MCMC variance across the entire range of *M* examined, and the shortfall does not vanish as *M* grows: the two converge to different limits (Table 2). This residual gap reflects the distinction between a frequentist asymptotic-likelihood approximation and a finite-sample Bayesian posterior. The GH_Logit_ estimate propagates the *asymptotic* (logit-normal) covariance of the maximum-likelihood estimates through the exact transformation, whereas the MCMC posterior reflects the actual finite-sample uncertainty, which the asymptotic approximation does not fully recover. Two alternative explanations can be set aside. Because we averaged over 100 independent runs of the sampler, the difference is not Monte Carlo noise; and because it is still present at the largest augmentation sizes examined, it is not an artifact of insufficient augmentation.

## 5 Considerations for practitioners

The preceding sections establish a single, practical lesson: because *M* is a prior and not merely a computational setting, it must be chosen with care. Two features of that choice are common to both inferential frameworks. First, there is a hard lower bound: if *M* is chosen too small – specifically *M* < *f*_0_ the finite binomial ceiling is not just a theoretical concern. Because the analytical variance relies on the curvature of the likelihood surface at the maximum-likelihood estimate, pinning the estimate against the arbitrary boundary at *N*_max_ = *D* + *M* produces an artificially flat surface and a suppressed variance (Section 4); the Bayesian posterior is suppressed in the same way, its upper tail truncated against the ceiling (Fig. 1). No choice of prior rescues *M* < *f*_0_. Second, once *M* > *f*_0_ the point estimate 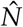 stabilizes, and the variance estimate climbs toward its target and then plateaus short of it, as the binomial converges to the Poisson distribution. Choosing *M* too aggressively on the low end is a recipe for overconfidence; the shared imperative is therefore to clear *f*_0_ comfortably.

Where the two frameworks diverge is in what happens above that lower bound. The divergence is computational in origin, but it has a statistical consequence: the cost of the sampler is what determines whether the choice of *M* can be sidestepped or must be chosen carefully. It might seem reasonable to simply make *M* as large as computationally tolerable. In a maximum-likelihood framework this instinct is largely correct: there is no statistical penalty for a generous *M*. The point estimate is invariant to *M* once *M* > *f*_0_, and the variance estimate is stable; bigger is, in a real sense, simply safer – though, as noted below, this depends on the parameters being well identified. The only qualification is numerical. As *M* grows, 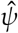 shrinks toward zero, and at extreme values it may approach the limits of numerical precision in some optimization software – though this threshold lies far beyond any realistic application.

For Bayesian implementations, the foremost consideration is computational: each augmented individual adds a latent binary variable for the sampler to update, so cost scales linearly with *M* – doubling *M* roughly doubles run time. A practitioner therefore cannot set *M* enormously large “to be safe” as one would in the likelihood framework; an extravagant *M* may render the sampler intractable. Finding an adequate value of *M* requires balancing computational feasibility against the requirement that *M* comfortably exceeds *f*_0_. Prior choice deserves a caveat here. In the well-identified *M*_0_ example examined above, a standard uniform prior on the inclusion probability *ψ* performs well, and the scale prior [*N*] *∝*1/ *N* advocated by Link (2013) returns practically the same answer. That equivalence should not be generalized too readily. Link’s own reanalysis of a heterogeneity model (*M*_*h*_) under the discrete uniform prior gave a posterior standard deviation for *N* of 20.0 at *M* = 168 and 67.6 at *M* = 1000: rather than plateauing once the ceiling stops binding, as it does in Table 2, the posterior simply kept expanding into whatever room the augmentation allowed. Where the parameters are weakly identified – sparse data, individual heterogeneity, or both – the advice to set *M* generously and stop worrying is therefore unsafe, and the scale prior is the more defensible choice. In the well-identified case, the key imperative is not prior choice but ensuring *M* > *f*_0_ with sufficient margin; once that condition is met, the remaining pressure to economize on *M* is purely computational. A practitioner who objects to the Binomial ceiling itself has a further option: the alternative specifications of Schofield & Barker (2014) place a Poisson prior on *N* directly, removing the Binomial variance constraint at any *M*, at the cost of a less standard model specification. This does not relax the lower bound — *N* remains bounded above by *M* — so clearing *f*_0_ comfortably is still required.

This also clarifies what diagnostics can and cannot tell you. Inspecting the posterior of *f*_0_ for mass accumulating near the augmentation ceiling *M* (Fig. 1) is the key check for the *lower*-bound failure: visible mass at the boundary signals that *M* is too small and must be increased. It does not, however, speak to the other end of the range: once *M* comfortably exceeds *f*_0_, the posterior of *f*_0_ looks equally well-behaved for any larger *M*, so the diagnostic confirms only that the lower bound has been cleared.

In short, the underlying statistical logic is identical across frameworks, but the practical imperative differs. In every case, clear *f*_0_ comfortably – that lower bound is non-negotiable. If you are working in a likelihood framework and computation is cheap, then make *M* large; the prior’s ceiling is lifted at no computational cost. If you are working in a Bayesian framework, the same lower bound applies but computation sets the ceiling: locating a value of *M* that clears *f*_0_ with margin is a genuine statistical investment, and above that the choice reduces to balancing adequacy against the linear cost of the sampler. Either way, *M* is part of model specification, not tuning.

## Summary

Data augmentation establishes a fundamental equivalence between two widely used modeling frameworks in quantitative ecology: occupancy modeling and closed-population capture–recapture. By augmenting an observed set of *D* capture histories with *M* all-zero histories and fitting a standard single-season occupancy model to the resulting dataset, one recovers maximum likelihood estimates of abundance that are numerically identical to those obtained from classical capture–recapture estimators. The occupancy parameter *ψ* in the augmented model corresponds to the inclusion probability, and abundance is estimated as 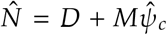, where *ψ*_*c*_ is the inclusion probability conditional on non-detection (equivalently, 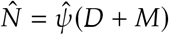 evaluated at the MLE). We develop this connection rigorously, derive analytical variance estimators for 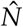 using the Delta method and a second-order Hessian correction, and demonstrate that Gauss-Hermite quadrature substantially narrows the gap between the local analytical approximations and MCMC-based estimates, leaving only a small residual. That gap arises from the asymmetric, heavy-tailed distribution of the conditional probability *ψ*_*c*_ and persists even once *M* is large enough for the binomial ceiling itself to stop binding. The central finding is that *M* is not merely a computational parameter but a hidden prior: it specifies a finite binomial ceiling that systematically constrains the variance of 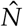. We provide framework-specific guidance on choosing *M*: in likelihood-based analyses, *M* should be set generously since computation is cheap; in Bayesian MCMC implementations, where computational cost scales linearly with *M*, locating a defensible lower bound on adequate augmentation size represents a statistically meaningful decision that warrants careful attention.

## 6 Acknowledgements

This manuscript benefited from the insights of several colleagues during its development. Larissa Bailey and Gary White (Colorado State University) provided suggestions on framing abundance estimation within an occupancy modeling context and insights into the computational details of Gauss-Hermite quadrature, respectively. Nathan Hostetter (USGS NC Cooperative Fish & Wildlife Research Unit) and Ben Augustine (USGS Rocky Mountain Science Center) provided constructive feedback that significantly improved the presentation and focus. We especially thank Ben Augustine for highlighting connections to previous work by Matt Schofield and Richard Barker. Any use of trade, firm, or product names is for descriptive purposes only and does not imply endorsement by the U.S. Government.

## Addendum 1: First-Order Delta Approximation for var(*ψ;*_*c*_)

From eqn. (2), let the transformation function *g* be the conditional inclusion probability:

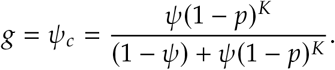

We approximate the variance of 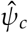 using the first-order Delta method as:

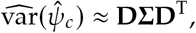

where *Σ* is the estimated covariance matrix of 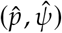 on the probability scale, and **D** is the Jacobian of *g* with respect to *p* and *ψ*:

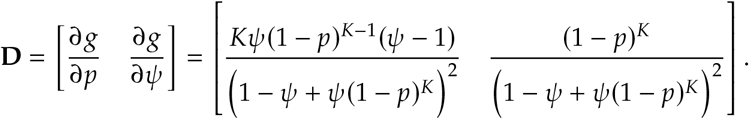

Applying this to the *Z* = 50 augmented dataset, where 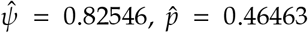 (such that *p*^*^ = 0.84655), *K* = 3, and *M*_*t*+1_ = 116. The empirical variance-covariance matrix for the probability-scale parameters is:

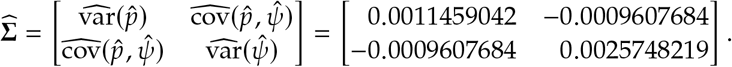

Evaluating the first-order approximation yields:

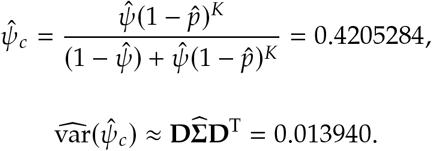

## Addendum 2: A second-order correction for var(*ψ;*_*c*_)

The first-order approximation of Addendum 1 treats the transformation *g* = *ψ*_*c*_ as locally linear, which artificially clips the right-skewed heavy tail produced when *M* is finite. Retaining the next term in the Taylor expansion recovers some of this omitted variance by incorporating the curvature of *g* through the Hessian matrix (**H**) of second partial derivatives:

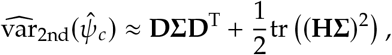

where **H** is defined as:

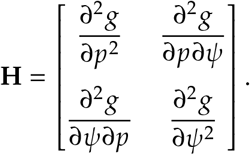

Let the denominator of our function *g* be *W* = 1 − *ψ* + *ψ*(1 − *p*)^*K*^. The second partial derivatives are:

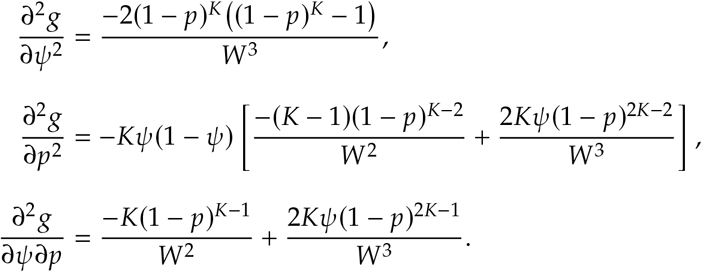

Returning to the *M* = 50 example 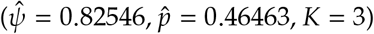, evaluating **H** and calculating the trace component yields a second-order correction of 0.000362. Adding this to the first-order estimate (0.013940) provides the revised conditional variance 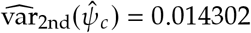.

Substituting this revised conditional variance into the strict variance estimator (eqn. 8) increases the total analytical variance for abundance 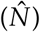 at *M* = 50 from 46.337 to 47.224. However, as Table 2 shows across a wide range of augmentation sizes, this correction recovers only a small part of the shortfall: the second-order term is still local curvature evaluated at a single point, whereas the unaccounted variance stems from the entire right tail.

## Addendum 3: Numerical Integration Using Gauss-Hermite Quadrature

Gauss–Hermite (GH) quadrature approximates an expectation under a Gaussian weight by a weighted sum over a fixed set of nodes. Applying it here requires first deciding the scale on which the two parameters – the detection probability *p* and the inclusion probability *ψ* – are to be treated as (approximately) Gaussian.

Because *p* and *ψ* are both confined to [0, 1] , it might seem reasonable to integrate on the probability scale, placing a Beta distribution on each parameter (with shape parameters set from the point estimate and its variance by the method of moments) so that no node can stray outside [0, 1]. Two things argue against it. First, the maximum-likelihood estimates were obtained by unconstrained optimization on the logit scale, so the inverted observed-information matrix is an asymptotic covariance for the *logit-scale* parameters: it describes a bivariate normal in logit space, not a pair of Betas in probability space. Second, a bivariate Beta has no convenient covariance structure (Olkin & Trikalinos 2015), so nodes built from two independent Betas discard the estimated correlation between *p* and *ψ*. The logit scale avoids both problems: it carries the full asymptotic covariance, including the correlation, it is the scale on which the normal approximation holds, and the inverse-logit transform returns every node to [0, 1] automatically.

Therefore, integration must occur on the unbounded logit scale. By matching the exact distributional assumptions of the asymptotic likelihood theory, we can evaluate the exact transformation function over the true probability mass. Let *θ* = [*α, β*]^T^, with *α* = logit(*p*) and *β* = logit(*ψ*). Unconstrained optimization returns the estimates 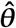 and the asymptotic covariance 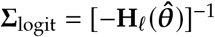, where **H**_*ℓ*_ is the Hessian of the log-likelihood.

We generate standard normal Gauss-Hermite quadrature nodes, **z**_*i*_, and associated weights, *w*_*i*_. These independent nodes are mapped into the correlated logit parameter space using the Cholesky decomposition of the covariance matrix (Press *et al*. 2007), **L** (where *Σ*_logit_ = **LL**^T^), via the affine transformation:

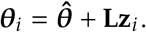

This produces a grid of discrete points [*α*_*i*_ , *β*_*i*_]^T^ that captures the multivariate normal approximation.

Each node is back-transformed to the probability scale:

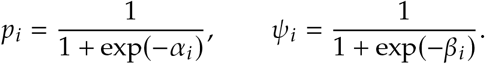

At each node, the conditional inclusion probability is evaluated exactly:

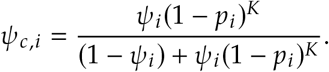

Applying the Law of Total Variance across the weighted nodes yields the variance of the abundance estimate,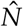. This computes both the expected binomial variance at each node and the variance of the expected population sizes across the distribution, which maps directly to the first-principles derivation in eqn. (6):

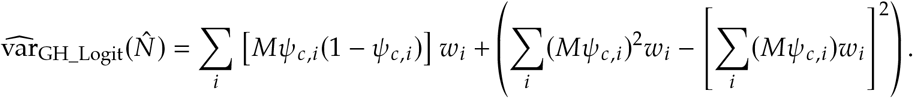

## Footnotes

1 We use *D*, rather than the more common *M*_*t* + 1_, to minimize confusion with our subsequent use of *M* to represent the size of the ‘augmented sample’

2 Following Otis *et al*. (1978), *M*_0_ denotes the constant-*p* closed-population model; this is unrelated to our use of *M* for the augmentation size.

